# Origin and functional diversification of PAS domain, a ubiquitous intracellular sensor

**DOI:** 10.1101/2023.04.23.537977

**Authors:** Jiawei Xing, Vadim M. Gumerov, Igor B. Zhulin

## Abstract

Signal perception is a key function in regulating biological activities and adapting to changing environments. PAS domains are ubiquitous sensors found in diverse receptors in bacteria, archaea, and eukaryotes, but their origins, distribution across the tree of life, and extent of their functional diversity remain unknown. Here, we show that using sequence conservation and structural information it is possible to propose specific and potential functions for a large portion of nearly 3 million PAS domains. Our analysis suggests that PAS domains originated in bacteria and were horizontally transferred to archaea and eukaryotes. We reveal that gas sensing via a heme cofactor evolved independently in several lineages, whereas redox and light sensing via FAD and FMN cofactors have the same origin. The close relatedness of human PAS domains to those in bacteria provides an opportunity for drug design by exploring potential natural ligands and cofactors for bacterial homologs.

**Teaser:** Signaling domains that originated in bacteria hold a potential as drug targets in humans.

## Introduction

Signal transduction pathways in all living cells detect nutrients, hormones, oxygen, redox potential, and other signals, and regulate various cellular functions accordingly. Signals are recognized by receptor proteins via dedicated sensor domains. PAS (**<u>P</u>**er-**<u>A</u>**rnt-**<u>S</u>**im) domains are ubiquitous sensor domains that are found in transcription factors, chemoreceptors, histidine and serine/threonine protein kinases and phosphatases, enzymes controlling second messenger turnover, ion channels, and other signaling proteins (*1*). While many sensor domains are extracytoplasmic, PAS domains are exclusively located in the cytoplasm (*1*). PAS domains adopt a conserved globular fold with a distinct binding cavity that can accommodate various ligands and cofactors thus enabling sensing capability (*2*). In addition to the sensory role, PAS domains may act as signal transducers and promote protein-protein interactions (*1*). PAS domain containing signaling pathways control complex behaviors ranging from motility (*3*), quorum sensing (*4*), and virulence (*5*) in bacteria to phototropism (*6*), circadian rhythms (*7*), oxygen homeostasis (*8*), and immune response (*9*) in eukaryotes. PAS domains in human transcription factors have become important drug targets for cancer therapy (*10–12*).

Over a million of PAS domain containing proteins are identifiable in current databases. However, signals recognized by these domains have been experimentally identified only in a handful of model proteins (Table S1). PAS domains are classified into several families based on sequence similarity, but the current classification does not reflect their biological functions. This is largely due to the extreme sequence divergence among PAS domains and the fact that only a few residues define their signal specificity. PAS domains are found in organisms ranging from bacteria to humans suggesting that they might have originated in the common ancestor. However, their evolutionary history, distribution across the Tree of Life, and potential functional diversification remain largely unknown.

Here we provide an extensive comparative genomic analysis of PAS domains across bacteria, archaea, and eukaryotes. We show that PAS domains originated in bacteria and evolved sensory functions through different paths. Eukaryotes acquired PAS domains from bacteria via many independent horizontal gene transfer events. Similarity of some of the human PAS domains to those in bacteria suggests that bacterial proteins can serve as useful models to determine signal specificity of the human counterparts, as it was recently shown for another ubiquitous sensor domain (*13*). Our findings open new avenues for functional studies and drug development using PAS domains, and our approach can serve as a framework for studying other protein families with extremely diverse members and versatile functions.

## Results

### PAS domain distribution across the tree of life

The distribution of PAS domains across the tree of life has not been systematically investigated since the original discovery of this superfamily, which involved a small number of genomes available at the time (*14, 15*). We searched for PAS domains in the entire set of UniProt reference proteomes (see Materials and Methods for details) and established that PAS domains are present in 66% of archaeal, 93% of bacterial, and 93% of eukaryotic proteomes (Data S1). In bacteria, PAS domains are widely present in most phyla; they are absent in the reduced genomes of endosymbionts and intracellular parasites, such as Dependentiae, Endomicrobia, Mycoplasmatales, and Rickettsiales (Fig. 1A, Data S1). In archaea, PAS domains have an uneven distribution: Halobacteriota have disproportionally more PAS domains than other phyla (Fig. 1B). In eukaryotes, PAS domains are widely distributed across different phyla except for Apicomplexa (Fig. 1C), which includes obligate endoparasites with reduced genomes, such as *Plasmodium*, *Babesia*, *Cryptosporidium*, and *Toxoplasma*.

**Figure 1.**
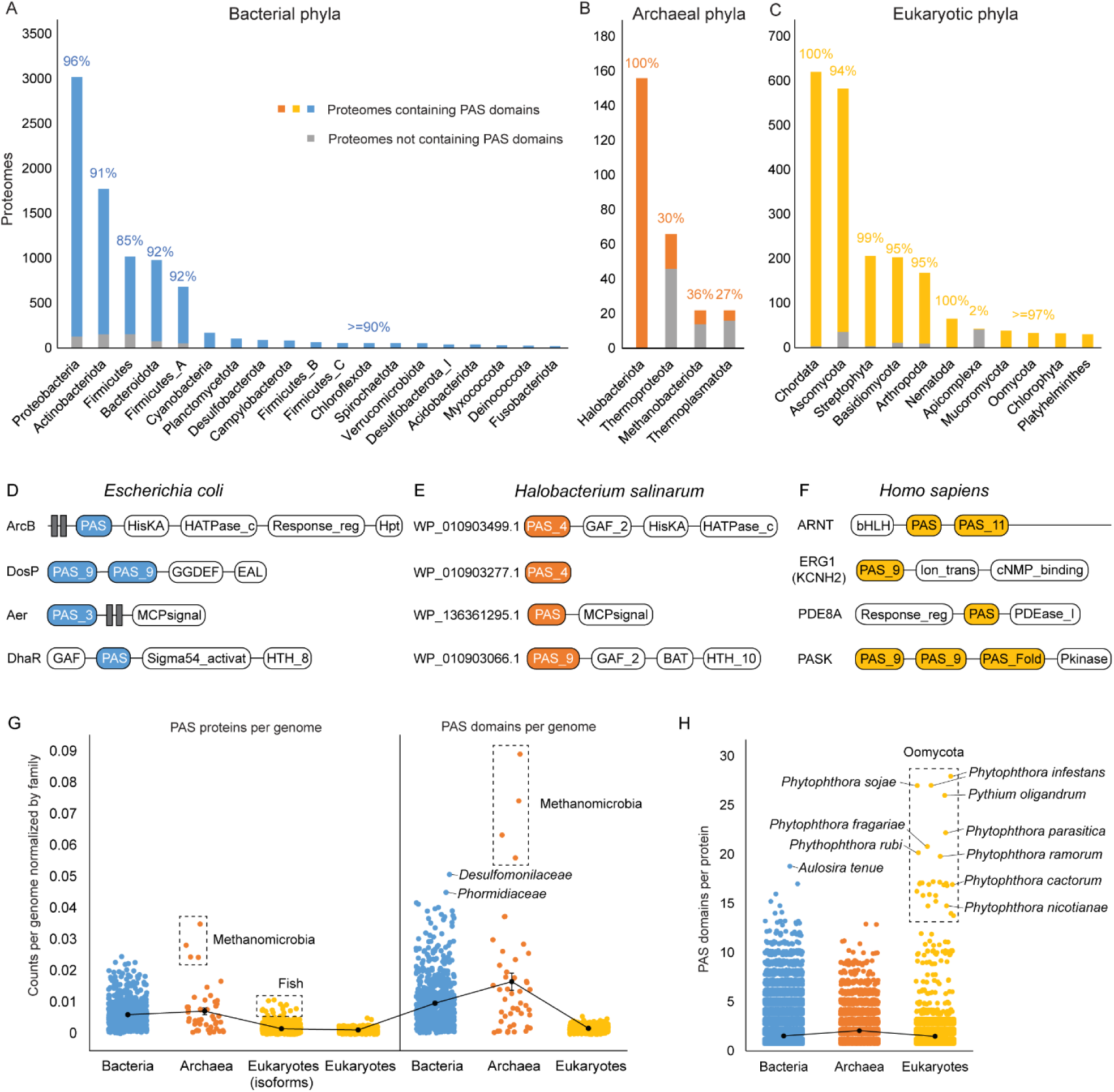
Global distribution of PAS domains. (**A-C**) PAS domain distributions in reference proteomes from bacterial (**A**), archeal (**B**), and eukaryotic (**C**) phyla. Colored bars show numbers of proteomes with PAS domains in each phylum. Grey bars show numbers of proteomes without PAS domains in each phylum. Percentages show proportions of PAS-containing proteomes in each phylum. Phyla with fewer than 20 reference proteomes are not shown. (**D**-**F**) Representative PAS-containing proteins in three model organisms. (**G**) Average numbers of PAS proteins per genome (left) and PAS domains per genome (right) across three kingdoms (normalized by total gene numbers, see Materials and Methods). Each dot represents the average count of a family. (**H**) Numbers of PAS domains per protein across three kingdoms. Each dot represents the number of PAS domains from a protein.

Next, we collected all PAS containing proteins defined by the Pfam PAS Fold profile (CL0183) from InterPro (*16*) and identified their domain composition (Data S2). We found that histidine kinases are the most common PAS proteins in bacteria and archaea, whereas transcription factors are more prevalent PAS proteins in eukaryotes (Fig. S1). Examples of typical domain architectures of the PAS proteins in three model organisms are shown in Fig. 1D-F.

To find out whether PAS domains and PAS containing proteins are enriched in some taxa, we quantified their content for each genome (normalized by the genome size at the family level, see Materials and Methods). Because of alternative splicing, we calculated the number of eukaryotic PAS proteins both with and without isoforms (Fig. 1G). We found that archaeal and bacterial genomes on average encode more PAS proteins and PAS domains and have higher variances than eukaryotic genomes (Fig. 1G). Moreover, all 17 Pfam PAS families are found in bacteria, whereas only 10 PAS families were found in eukaryotes and 7 in archaea (Table S2). Within each kingdom, the number of PAS proteins and PAS domains vary in different taxonomic groups (Data S2). For example, genomes of *Desulfomonilaceae* (desulfobacteria) and *Phormidiaceae* (cyanobacteria) are enriched with PAS domains (Fig. 1G, blue). In archaea, the Methanomicrobia class (Halobacteriota) encodes a substantially higher proportion of PAS proteins and PAS domains (Fig. 1G, orange). In eukaryotes, teleost fishes have the highest number of PAS proteins especially when considering isoforms (Fig. 1G, gold).

On average, PAS proteins in all three kingdoms contain two PAS domains (Fig. 1H). However, some organisms have proteins with a much larger number of PAS domains. For example, *Aulosira tenue* and some other cyanobacteria contain up to 19 PAS domains in a single histidine kinase (Fig. 1H, blue; UniProt: A0A1Z4MZL8). Remarkably, histidine kinases with more than 14 PAS domains are also found in eukaryotes, specifically, in notorious plant pathogens of *Phytophthora* genus (Fig. 1H, gold; Data S2). We then analyzed the relationship between the number of PAS domains and the genome size and found statistically significant correlation between the number of genes and the number of PAS proteins and PAS domain (Fig. S2; see Materials and Methods).

We noticed that some of eukaryotic PAS domains were not identified by Pfam profile models and the default search tool HMMER (e.g., the third PAS domain of human PASK in Fig. 1F), indicating that the actual number of eukaryotic PAS domains may be higher. To explore this, we compared the number of PAS domains identified in 14 representative eukaryote genomes by HMMER (*17*) with that identified by a manually curated structure search FoldSeek (*18*). Strikingly, 36.4% PAS domains revealed by FoldSeek were not identified by HMMER, indicating that the thresholds used by Pfam were too strict (Data S3). Therefore, we applied a released threshold for HMMER (E-value < 1) and collected PAS proteins from 43 representative eukaryotic genomes. False positive matches without a PAS fold have been removed by manually checking the ESMFold structures (*19*). As a result, we found that most eukaryotes have around 10-30 PAS proteins per proteome, but much larger numbers (50-80 PAS proteins) are seen in *Naegleria gruberi*, *Acanthamoeba castellanii*, *Guillardia theta*, and *Danio rerio* (Data S4). Moreover, alternative splicing in Chordata results in up to 300-400 PAS isoforms per proteome (Data S4).

### Current classification of PAS domains does not reflect their biological functions

To understand whether the current classification of PAS domains reflects their biological functions, we conducted a comprehensive literature search for cofactor-binding PAS domains and compared their distribution within Pfam protein families (*20*) (Table S1). We established that the current classification of PAS domains does not reflect sensory functions: (i) PAS domains with the same cofactor can be found in different families and (ii) the same family can contain PAS domains with different cofactors (Table S1). Furthermore, we found that profile models for two large families are incorrect: PAS_3 (Pfam entry: PF08447) does not include the N-terminal Aß and Bß, and PAS_8 (Pfam entry: PF13188) does not include the C-terminal Gß, Hß, and Iß (*20*) (Fig. S3A, Table S3). Next, we performed all-vs-all BLAST searches using seed sequences from all families after adding missing regions to PAS_3 and PAS_8. We found that while small families form distinct clusters, the large families, including PAS, PAS_3, PAS_4, PAS_8, and PAS_9 (Pfam entries: PF00989, PF08447, PF08448, PF13188, PF13426), are mixed in the sequence similarity network, indicating overlaps and misclassification (Fig. S3B).

### Functional PAS families can be identified using sequence and structural information

Using HMMER, we collected approximately 2.8 million sequences that belong to 17 PAS families from the RefSeq database (*21*) and searched sequences of each PAS family against the entire Pfam database (*20*) (see Materials and Methods). Noticeably, we observed numerous cases of overlaps between nine major families, including PAS, PAS_3, PAS_4, PAS_7, PAS_8, PAS_9, PAS_10, PAS_11, and PAS_12 (Fig. S4, Table S4). According to the InterPro database, sequences of these families comprise more than 80% of the entire PAS fold superfamily (*16*). To reclassify these families, we combined these sequences and reduced redundancy over 80% identities, which resulted in 369,910 PAS domain sequences (Data S5). Next, we performed an all-versus-all sequence alignment using DIAMOND (*22*) and assigned sequences into MCL clusters based on the sequence similarity network (*23*). This resulted in 462 clusters, 163 of which contain at least 100 sequences and comprised 98% of the entire dataset (Fig. 2A, S5, Data S6). The size of the cluster correlates with the average sequence length of its members (Fig. S5), indicating that clusters 164-462 may consist of partial sequences. Out of the 163 clusters, more than 100 (over 300,000 sequences) have conserved residues inside the PAS domain cavity that might serve as potential binding sites for cofactors or ligands (Fig. 2A; all data available at https://github.com/bioliners/PAS).

**Figure 2.**
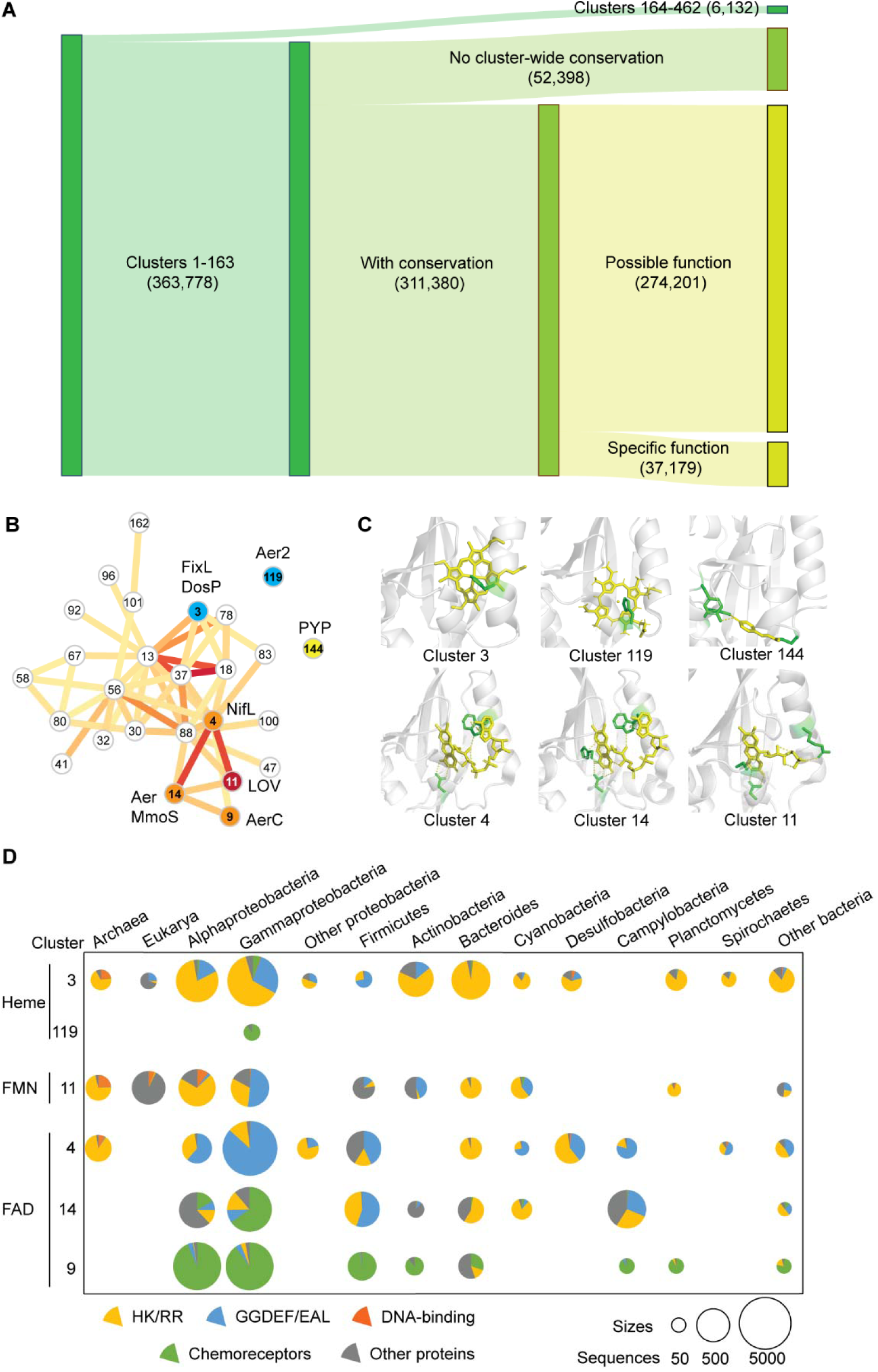
Functional clusters of PAS domains. (**A**) Clustering results. Possible functions stand for sequences with conservation but unknown functions. Specific functions indicate sequences with predicted cofactors. Numbers of sequences in each category are shown. (**B**) Markov clusters of PAS domains. Clusters are numbered according to the number of sequences (clusters with smaller numbers contain more sequences). Colored edges connecting clusters reflect BLAST hits between clusters (yellow to red from few to many). Only clusters 1-163 with conserved residues within the cavity are shown. Outlier clusters are not shown for simplicity. Clusters with specific conserved cofactors are highlighted: blue, heme-binding; orange, FAD-binding; red, FMN-binding; yellow, PYP homologs. Representative proteins are labeled next to these clusters. (**C**) Cofactor-binding structures. A representative PAS domain is shown for each cluster: cluster 3, FixL (PDB: 1DRM); cluster 119, Aer2 (PDB: 4HI4); cluster 144, PYP (2PHY); cluster 4, NifL (PDB: 2GJ3); cluster 14, Aer (PDB: 8DIK); cluster 11, Phot1 (PDB: 2Z6C). Yellow, cofactor; green, conserved residues for cofactor-binding. (**D**) Distribution of PAS domain clusters. The size of the chart represents the number of PAS domains. Colors indicate PAS domains from different proteins. HK/RR, two component systems; GGDEF/EAL, cGMP regulators.

Some of these clusters contain well-studied PAS domains with known cofactors (Fig. 2B, Table S1). For example, cluster 3 includes PAS domains of the histidine kinase FixL and the diguanylate cyclase DosP, both of which have a histidine at Fα serving as a heme-binding site (*24, 25*) (Fig. 2C, Table S1). In more than 9,700 (60%) sequences in this cluster this histidine is conserved, strongly suggesting that all these homologs bind heme as the cofactor (Fig. S6). Custer 119 contains PAS domains from the aerotaxis receptor Aer2, which has a histidine for heme-binding at a different location, Eα (Eη) (*26*) (Fig. 2C, Table S1). More than 200 (73%) sequences in cluster 119 have this conserved histidine, suggesting heme-binding for all these homologs (Fig. S6). Clusters 4, 9, and 14 contain PAS domains of the histidine kinases NifL and MmoS, and the chemoreceptors AerC and Aer, all of which are redox sensors with FAD as the cofactor (*27–30*) (Fig. 2C, Table S1). A conserved tryptophane at Fα was identified as the only key residue for FAD-binding (*30*). Across the three clusters, more than 22,600 sequences (53%, 91%, and 84%, respectively) have tryptophane in the same position, suggesting FAD as the cofactor for all these PAS domains (Fig. S6). Cluster 11 contains a PAS domain subfamily called LOV (Light-Oxygen-Voltage) domains, which serve as blue light sensors and have a conserved cysteine for covalent bonding to FMN, FAD, or riboflavin (*31–33*) (Fig. 2C, Table S1). More than 4,300 (57%) PAS domains from this cluster have the conserved cysteine, suggesting binding of these cofactors (Fig. S6). Cluster 144 contains a different blue light sensor – photoactive yellow protein (PYP), which has conserved cysteine, tyrosine, and glutamate for *p*-coumaric acid binding (*34*) (Fig. 2C, Table S1). The three residues are also conserved in this cluster, indicating *p*-coumaric acid as their cofactor (Fig. S6).

Our clustering approach enabled assignment of specific cofactor binding and indication of potential ligand/cofactor binding to more than 80% of PAS domains with currently unknown functions (Fig. 2A). It further confirmed that PAS domains have a broad phyletic distribution, especially in bacteria (Fig. 2D, Data S7). Furthermore, some clusters are enriched in distinct signal transduction proteins. For example, clusters 3 (heme), 4 (FAD), and 11 (FMN) are associated with histidine kinases and diguanylate cyclases, clusters 9 (FAD) and 119 (heme) with chemoreceptors, and cluster 14 (FAD) with all three types of receptors (Fig. 2D, Data S7).

### Heme and flavin containing PAS domains have different evolutionary paths

PAS domains have short and diverse sequences, which complicates their phylogenetic analysis (*1*). However, they all share the same structural fold (*35*). Here we sampled 1% sequences from each cluster and constructed phylogenetic trees based on a previously established structure-guided approach designed for short and diverse sequences (*36, 37*) (see Materials and Methods). We found that clusters 3 and 119 that contain known heme-binding PAS domains are not related to each other (Fig. 3A). Other putative heme-binding PAS domains are also found in different clusters (Table S1): specifically, ERG3(KCNH7)-PAS (*38*) is in cluster 11, CLOCK/NPAS2-PAS B (*39, 40*) are in cluster 72, and CLOCK/NPAS2-PAS A (*39, 40*) are in cluster 87, all of which are also not related to clusters 3 and 119 (Fig. 3A). In addition, the histidine residue for heme-binding is also located in a different position in each cluster (*38, 39*) (Fig. 2C, 3B). Taken together, the sequence, structural, and phylogenetic information suggests that PAS domains in these clusters evolved heme-binding capabilities independently (Fig. 3C).

**Figure 3.**
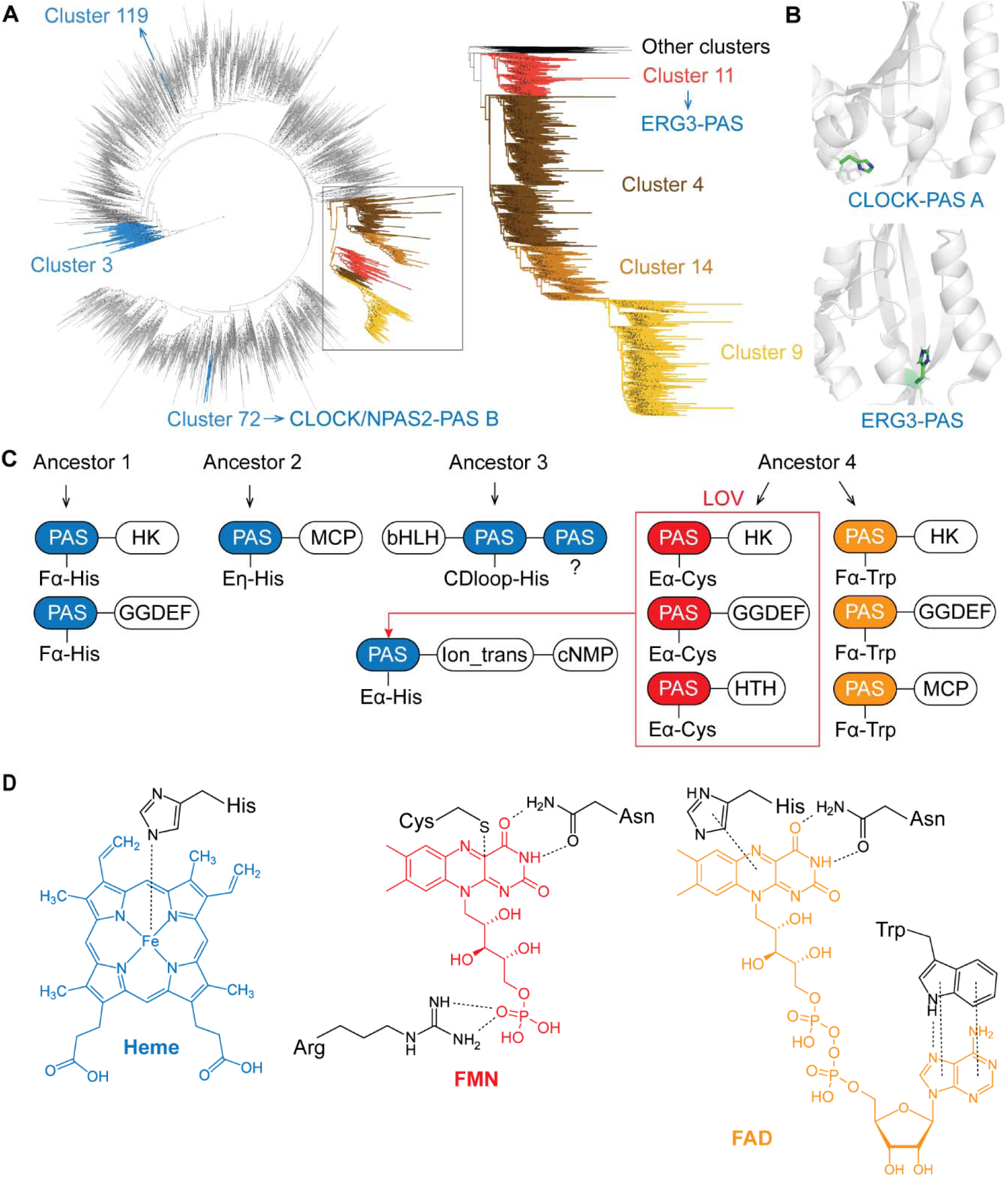
Evolutionary history of cofactor-binding PAS domains. (**A**) Maximum likelihood tree using sampled sequences from each cluster. Clusters of interest are marked with different colors. Clusters 4, 9, 11, and 14 form a single clade, and a more accurate tree is shown on the right using all sequences from these clusters after reducing 60 % sequence redundancy. Dots on the tree indicate bootstrap values not less than 70. (**B**) Structures of heme-binding PAS domains. ERG3-PAS (PDB: 6Y7Q) and CLOCK-PAS A (PDB: 6QPJ) are shown. Putative residues for heme coordination are highlighted in green. ERG3-PAS is from cluster 11 (shown in panel A). CLOCK/NPAS2-PAS A is from cluster 87 (not included for tree building due to low conservation). CLOCK/NPAS2-PAS B is from cluster 72 (shown in panel A) and may also bind heme, but the putative binding site is unknown. (**C**) Evolutionary paths of heme and flavin binding PAS domains. Ancestor 1, ancestor of cluster 3 PAS domains; Ancestor 2, ancestor of cluster 119 PAS domains; Ancestor 3, ancestor of bHLH-PAS; Ancestor 4, common ancestor of clusters 4, 9, 11, and 14. Blue, heme-binding PAS domains; red, FMN-binding PAS domains; orange, FAD-binding PAS domains. Key residues for cofactor-binding are labeled for each PAS domain. Light-Oxygen-Voltage (LOV) domains bind FMN (or FAD) for blue light sensing. (**D**) Scheme of heme and flavin binding sites. Key residues for cofactor-binding are shown.

In contrast, clusters 4 (FAD), 9 (FAD), 11 (FMN), and 14 (FAD) form a single clade on the tree (Fig. 3A). We also built a more accurate tree using all sequences from these four clusters (after reducing redundancy) and found that cluster 11 (FMN) and other clusters (FAD) form two branches, indicating functional divergence from a single origin (Fig. 3A, right). Furthermore, a similar cofactor-binding site is present in all four clusters (Fig. 2C). Taken together, these data suggest that FMN and FAD binding PAS domains evolved from the same origin, and then diverged into four clusters (Fig. 3C). Intriguingly, the heme-binding PAS domain from ERG3 (KCNH7) is a homolog of cluster 11 LOV domains, indicating neofunctionalization from FMN to heme binding (Fig. 3C). A scheme for binding of the three cofactors in PAS domains is shown in Fig. 3D.

### Bacterial origins of eukaryotic PAS domains

Using the sequence and structure searches, we identified 34 PAS proteins in the human genome (Data S3, S4, S8; see Materials and Methods). A literature search showed that these proteins play important roles in human health (Table S5). Based on the domain architecture, these proteins can be grouped into four families: the potassium ion channels KCNH, the phosphodiesterases PDE8, the transcription factors bHLH-PAS, and the serine/threonine kinase PASK (Fig. 1F, 4, Table S5). According to the analysis of our representative genome dataset, all these four families were present prior to the emergence of Metazoa, however, the PAS domains were integrated into these proteins during early stages of metazoan life (Fig. 4A, Data S4). To investigate origins of these PAS domains, we performed BLAST searches against the NCBI RefSeq database using human PAS orthologs from the organisms located closest to the root of the tree (Fig. 4A): Remarkably, these searches identified no similar sequences in eukaryotes other than Metazoa but found PAS domains from various bacterial proteins with more than 40% sequence identity, suggesting their origins in bacteria (Data S9). To determine if this observation holds true for other eukaryotic PAS domains, we performed the same type of searches for randomly selected PAS domains from eukaryotes representing major lineages other than Metazoa: fungus *Spizellomyces punctatus* (RefSeq accession XP_016607844.1), choanoflagellate *Monosiga brevicollis* (XP_001742645.1), plant *Arabidopsis thaliana* (NP_001077788.1), cryptophyte *Guillardia theta* (XP_005835918.1), oomycete *Phytophthora sojae* (XP_009526312.1), alveolate *Perkinsus marinus* (XP_002767112.1), and excavate *Naegleria gruberi* (XP_002678807.1). Strikingly, we observed the same pattern: in each case, no eukaryotic sequences from other lineages were found, but instead diverse bacterial PAS domains with 35% - 48% sequence identities were identified (Data S9). Taken together, these data suggest that eukaryotes acquired PAS domains from bacteria via multiple independent horizontal transfer events.

**Figure 4.**
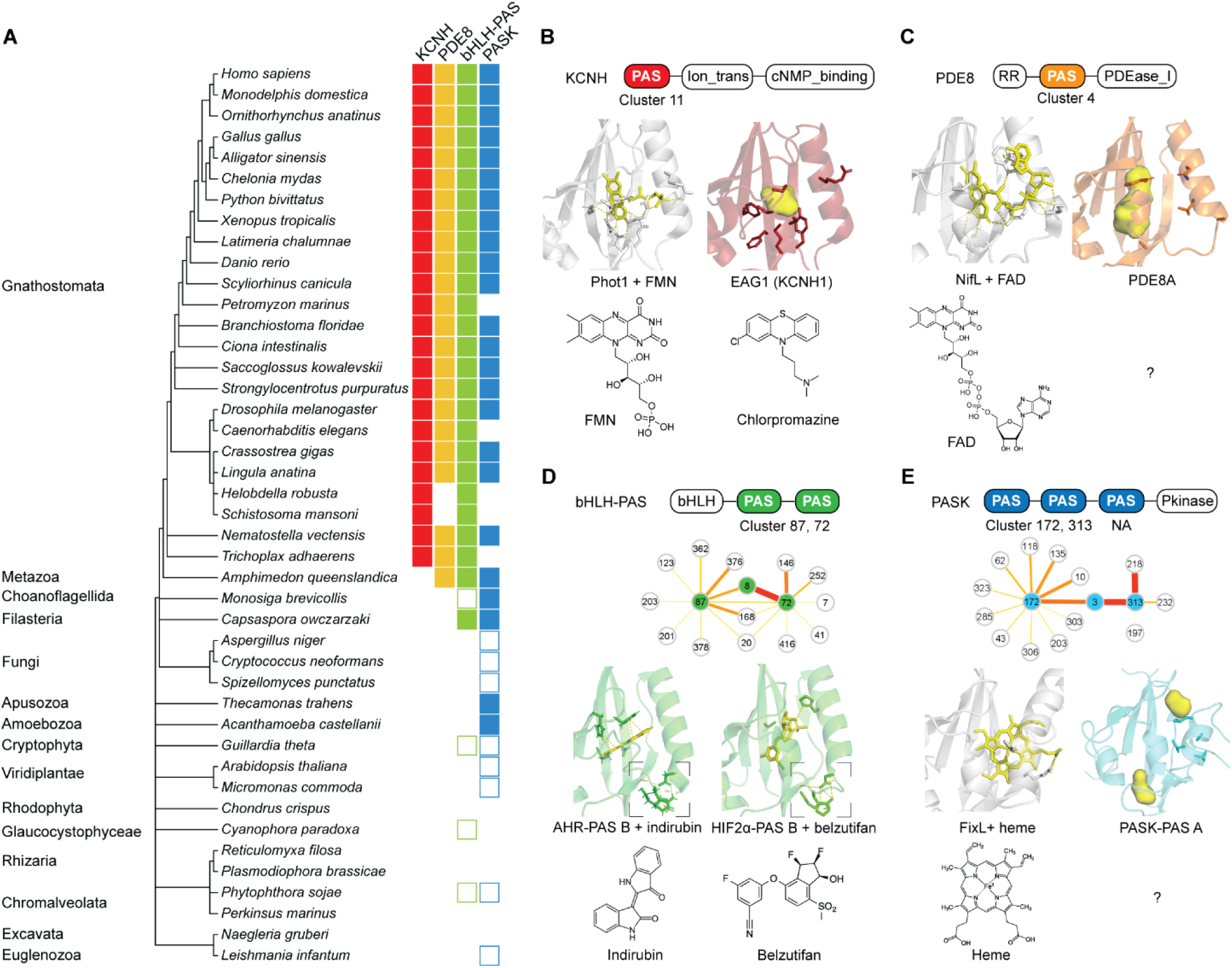
Evolutionary origin of human PAS domains. (**A**) Distribution of human PAS protein homologs across the tree of life. The tree of life is modified based on NCBI taxonomy browser. Filled boxes show organisms with PAS proteins homologous to human proteins. Empty boxes show organisms with similar PAS proteins but not homologous to human proteins (e.g., the signaling domain is a different version). (**B**) Potassium ion channel KCNH (e.g., EAG1, ERG3). KCNH-PAS belongs to cluster 11 (FMN). The grey structure shows the FMN-binding PAS domain of *A. thaliana* phototropin 1 (PDB: 2Z6C). The red structure shows the mouse EAG1-PAS with a putative drug-binding site (PDB: 4HOI). The drug chlorpromazine is a flavin analog. (**C**) Phosphodiesterase PDE8 (including PDE8A and PDE8B). PDE8-PAS belongs to cluster 4 (FAD). The grey structure shows the FAD-binding PAS domain of NifL (PDB: 2GJ3). The orange structure shows the human PDE8A-PAS with a putative binding site (AlphaFold). (**D**) bHLH-PAS transcription factors (e.g., CLOCK, NPAS2, AHR, HIF2α). PAS domains form their own clusters: PAS A form cluster 87 and PAS B form cluster 72. Both clusters are close to cluster 8 (clusters with average BLAST hit > 0.001 are shown). Green structures show the ligand-bound PAS B of AHR and HIF2α (PDB: 7ZUB, 7W80). Dashed boxes indicate the conserved H(P)(D/E)D…R motif. This motif is also conserved in cluster 8 (structure model available at https://github.com/bioliners/PAS). (**E**) PAS domains of the serine/threonine kinase PASK. PASK-PAS from their own clusters: PAS A form cluster 172, PAS B form cluster 313, and PAS C does not match to any Pfam HMMs. Both clusters are close to cluster 3 (clusters with average BLAST hit > 0.001 are shown). The grey structure shows the heme-binding PAS domain of FixL (PDB: 1DRM). The blue structure shows the human PASK-PAS A with cavities (PDB: 1LL8).

### Human PAS domains as potential drug targets

To further investigate the bacterial origins of human PAS domains, we integrated information from clustering and phylogenetic trees. We found that KCNH-PAS belong to cluster 11 (FMN) and may originate from Alphaproteobacteria FMN-PAS, as previously suggested (*41*) (Fig. 4B, S7). Structure comparison reveals a putative binding site in EAG1(KCNH1)-PAS, where polar residues for flavin-binding are replaced by hydrophobic residues (Fig. 4B). Notably, the antipsychotic drug chlorpromazine, which is a hydrophobic flavin analog, binds EAG1-PAS and modulate channel activities (*42*) (Fig. 4B). Thus, EAG1-PAS may bind chlorpromazine in a similar way as bacterial PAS domains bind flavins. In addition, we found that PDE8-PAS belongs to cluster 4 (FAD) (Fig. 4C). Structure comparison also shows a putative binding site and hydrophobic residues, suggesting that PDE8-PAS might bind flavin analogs (Fig. 4C).

The two PAS domains of bHLH transcription factors form their own clusters (87 for PAS A and 72 for PAS B) (Fig. 4D). Cluster 87 has low sequence conservation, but cluster 72 groups with cluster 8 on the phylogenetic tree, suggesting that the bHLH-PAS may originate from cluster 8, most likely from the PAS domains in Firmicutes (Bacilli) (Fig. S8). Notably, both bHLH-PAS and cluster 8 have a conserved H(P)(D/E)D…R motif connecting the α-helix and ß-strand, but how this conserved structure affects PAS domain properties remains unknown (Fig. 4D, S6). Although cluster 8 is largely unstudied, ligand binding has been reported in the bHLH-PAS protein AHR and anticancer drug-binding in HIF2α (Fig. 4D) (*43, 44*). PAS domains in PASK also form their own clusters, with cluster 3 (heme), which primarily consists of bacterial sequences, being the closest neighbor (Fig. 4E). Moreover, PASK-PAS from *Acanthamoeba castellanii*, *Thecamonas trahens*, and *Capsaspora owczarzaki* are found in cluster 3, further suggesting their origins from bacterial heme-PAS (Data S3). However, in human PASK-PAS A the heme-binding residue is not conserved, and potential ligand-binding cavities are in a different location compared to bacterial heme-PAS, indicating that it evolved a different function (Fig. 4E).

## Discussion

The last time genome-wide survey of PAS domains was carried out in 1999, when only 300 PAS domains from fewer than 20 genomes were available for analysis (*1*). In this study, we analyzed nearly 3 million PAS domains from more than 100,000 genomes of bacteria, archaea, and eukaryotes. Here, we present new findings about evolution, genomic landscape, and functional diversification of these remarkable biological sensors. We now show that PAS domains are present in most life forms; they are absent from organisms with reduced genomes, such as endosymbionts and intracellular parasites (*45*). We found that the number of PAS domains per genome correlates with the genome size (Fig. S2), thus following the trend established previously for major signal transduction families in bacteria (*46*). PAS domains were identified in major types of signal transduction proteins. In bacteria, they are predominantly found in histidine kinases, whereas in eukaryotes they are more often found in transcription regulators (Fig. S1).

We demonstrate that the current classification of PAS domains and profile models used for their identification in genomic datasets are outdated and lack biological information. We showed that using advanced clustering methods, sequence conservation, and structural information, it is possible to produce a better classification system. Still, deriving PAS domain function from sequence and structure remains a grand challenge. First, sequences from the same cluster might have different functions due to recent neofunctionalization, as we have recently shown for the photoactive yellow protein (*47*). Here, we found similar cases. For example, most sequences in cluster 3 contain a conserved histidine, which is a known heme-binding site, but some sequences in the cluster have a different conservation pattern, resulting in association with a different cofactor, Fe-S cluster (*47*) (Table S1). Second, proteins from different clusters may have the same function – what is known as non-homologous, isofunctional proteins, or results of convergent evolution. Several examples of heme-binding PAS domains fall into this category. Third, homologous PAS domains with the same function may be found in different clusters. This could be due to their association with different downstream signaling domains. For example, FAD-binding PAS domains form three clusters (4, 9, and 14), but they are more related to each other than to any other cluster and share the same cofactor-binding site, suggesting the same function despite significant sequence divergence (Fig. 2B, 2C). A previous study identified two clusters of FAD-PAS chemoreceptors exemplified by Aer and AerC that have different domain composition (*30*). Satisfactorily, these clusters correspond to our clusters 14 and 9. In addition, here we identified a novel cluster (cluster 4), where PAS domains come from signaling proteins other than chemoreceptors.

Despite these problems, we were able to propose a specific function (a conserved binding site for a specific cofactor) or a potential function (a conserved pattern within a binding cavity, indicating potential cofactor or ligand binding) for a large portion of the vastly unstudied “galaxy” of PAS domains (Fig. 2A). The latter category involves ligand-binding PAS domains (*48*), that were not analyzed in this study, because of limited published data. Furthermore, we hypothesize that clusters of PAS domains without conspicuous conservation may perform other biological functions, such as promoting protein-protein interactions and serving as signal transducers, which does not require strict conservation of individual amino acid residues. Thus, our findings provide new opportunities for targeted biochemical characterization of this important, but largely unstudied superfamily.

Our study also offers a new view on the origin and evolution of PAS domains. The fact that PAS domains were identified in such diverse organisms as bacteria, archaea, and eukaryotes, including humans (*1*) suggested early origins of PAS domains, perhaps even their presence in the last universal common ancestor. Here we show that this is not the case. Considering our current understanding of the relationship between bacteria, archaea, and eukaryotes (*49*), several lines of evidence suggest that PAS domains originated in bacteria and were horizontally transferred to archaea and eukaryotes: (i) Bacteria have the widest phyletic distribution of PAS domains and nearly all bacterial phyla have a similar proportion of PAS domains per genome (Fig. 1A). In contrast, many archaeal phyla do not have any representatives with PAS domains. Among archaea, Halobacteriota have disproportionally more PAS domains than other phyla (Fig. 1B), which is consistent with a previous finding that they acquired many signal transduction genes from bacteria (*50*). (ii) Bacteria have the widest diversity of PAS domains: they come from all currently known PAS families, whereas many of these families are not present in archaea and eukaryotes (Table S2). (iii) Histidine kinases are the class of signal transduction proteins that are mostly enriched in PAS domains (Fig. S1). It was previously shown that histidine kinases likely originated in bacteria and were horizontally transferred to archaea and eukaryotes (*51*). (iv) Metazoan PAS domains are more similar to bacterial PAS domains than to those from other eukaryotic lineages, indicating horizontal gene transfer (Data S9). (v) Randomly selected PAS domains from major eukaryotic supergroups are more similar to bacterial PAS domains than to those from other eukaryotic supergroups, indicating horizontal gene transfer (Data S9). A nearly equal distribution of PAS domains among bacterial phyla suggests that the PAS domain emerged early in bacterial evolution.

With respect to the evolution of a specific function, our results suggest that while heme-PAS domains have evolved several times, flavin-PAS domains have the same evolutionary origin. Heme-binding usually requires only one conserved residue, a histidine, for iron-coordination, which may facilitate its *de novo* evolution (Fig. 2D). In contrast, flavin-binding is much more complex, as it requires multiple hydrogen bonds and stacking interactions, thus constraining *de novo* evolution (Fig. 2D).

Our finding that PAS domains in humans have close homologs in bacteria is exciting. An anticancer drug targeting the PAS B domain in the transcription factor HIF2α was recently approved by FDA (*12, 44*), revealing a promising role of PAS domains as drug targets (Fig. 4D). The bacterial origin of human PAS domains could provide valuable information for drug design. For example, we found that bacterial flavin-PAS is the origin of human KCNH-PAS and PDE8-PAS, both are potential targets for cancer therapies (*52, 53*). EAG1(KCNH1)-PAS is regulated by the antipsychotic drug chlorpromazine, which is one of the flavin analogs blocking the flavokinase activity (*42, 54*). The fact that EAG1-PAS evolved from flavin-PAS and binds a flavin analog suggests that potentially drugs for PAS domains can be designed based on their original sensory functions in bacterial homologs. This hypothesis is further supported by a recent finding that the Cache domain (a distant homolog of PAS domains) (*55*) in the human α2δ-1 protein binds GABA-derived drugs the same way as bacterial Cache domains bind their natural ligands – GABA and other amino acids (*13, 56*).

This study highlights key properties of PAS domains as universal molecular sensors – their plasticity, evolvability, and transmissibility. It also exposes the need for a better classification system and current limitations in their identification and function prediction. Conserved clusters of PAS domains with proposed functions present an opportunity to design targeted experiments for validation and further exploration of their diverse properties.

## Materials and Methods

### Data collection

PAS-containing Proteomes and corresponding genome assembly identifiers were retrieved from the InterPro database (*16*) using its API (PAS Fold, CL0183). We assigned the NCBI (*57*) and GTDB (*58*) taxonomy using taxonomy IDs and a custom python script. Protein and gene identifiers of PAS proteins were retrieved from the UniProt database (*59*). In the case of eukaryotes, the longest isoform of the PAS proteins was kept for the subsequent analysis. Protein domain compositions were retrieved from the UniProt database based on the available Pfam models (*20*). PAS protein counts and PAS domain counts were calculated using a custom python script. PAS protein and domain counts were normalized based on the total gene count of corresponding genomes. All scripts used are available at https://github.com/bioliners/PAS.

Hidden Markov Models (HMMs) of PAS domain families were downloaded from the Pfam database (*20*). These HMMs were used as queries in *hmmsearch* to collect PAS domain sequences from the NCBI reference sequence protein database (RefSeq) (*60*) with a threshold of E-value < 0.01 (*17*). To find overlaps between PAS domain families, these sequences were then searched against all HMMs from Pfam using *hmmscan* (*17*).

### PAS domain distribution analyses

To mitigate the issue of uneven distribution of sequenced genomes across taxonomic groups when assessing the distribution of PAS domains across the tree of life, we sequentially averaged normalized PAS protein and domain counts from the species level up to the family level and expressed the final values as a percentage. In this way even if bacteria have significantly greater number of orders per family (and correspondingly genera per order and species per genus), it gets compressed to a single value per family. At the family level bacteria and eukaryotes have the most comparable numbers of members (families) without compromising the power of statistical tests. Spearman test was used to investigate the correlation of the number of PAS proteins and PAS domains with genome size expressed as the number of all encoded genes in the analyzed genomes. The number of PAS proteins were subtracted from the total number of genes prior to calculations to alleviate possible self-correlation. Scripts used are available at https://github.com/bioliners/PAS.

### Sequence clustering

PAS domain sequences collected from RefSeq were combined and sequence redundancy was reduced using CD-HIT with a threshold of identify > 80% (*61*). A sequence similarity network was generated by DIAMOND in the sensitive mode (E-value < 0.05, query coverage > 80%) (*22*). The network was then used for clustering by the high-performance Markov cluster algorithm (HipMCL) with an inflation of 1.4 (*23, 62*). The distance between two clusters was calculated as the average number of DIAMOND hits, i.e., the total number of DIAMOND hits between them divided by the product of the numbers of sequences in the two clusters.

### Cluster analyses

For each clusters multiple sequence alignments were constructed using MAFFT with default parameters (*63*). Sequence logos were generated using WebLogo 3 (*64*) and Skyalign (*65*). For clusters 1-163 with conserved residues that may serve as binding sites, a representative sequence was selected, defined by the best hit in *hmmsearch* when searching sequences against the HMM of the cluster built by *hmmbuild* (*17*). The conserved residues were highlighted in the AlphaFold model of the representative PAS domain (*66*). For the clusters with putative cofactors, taxonomy was obtained from GTDB (for bacteria and archaea) (*58*) and NCBI (for eukaryotes) (*57*), and protein architectures were identified using TREND with default parameters (*67, 68*). Sequences, logos, and structure models for clusters are available at https://github.com/bioliners/PAS.

### Phylogenetic analyses

Approaches for structure-guided alignment and phylogenetic tree construction were modified from previous studies (*36, 37*). PDB structures (1DP6, 1NWZ, 1V9Z, 2GJ3, 2MWG, 2PD7, 2Z6C, 3BWL, 3EWK, 3FG8, 3K3C, 4F3L, 4HI4, 4MN5, 4ZP4, 5HWT, 6CEQ, 6DGA, 6KJU, 6QPJ) were used for structure alignment using MUSTANG (*69*). This alignment resulted in an HMM using *hmmbuild*, which guided the alignment of representative sequences from each cluster using *hmmalign* (*17*). After trimming gaps (gaps > 90%) and removing sequences with low coverages (query coverage < 80%), the alignment was used for building a maximum likelihood tree using IQ-Tree (*70*) with the LG+G4 substitution model proposed by ProtTest 3 (*71*) and 1000 bootstraps. 1% sequences were randomly sampled from each cluster using a custom Python script, from which a final sequence alignment was built based on the previous alignment and tree using SEPP (*72*). A final maximum likelihood tree was built from this alignment using IQ-Tree with the same parameters (*70*).

### Structure search in human proteomes

PAS B from BMAL1 (PDB: 2KDK) was identified as the representative human PAS domain by *hmmsearch* against HMM (*17*). This PAS domain was used to search against the human proteome (UP000005640) from the AlphaFold database (*73*) using TM-align with the default parameters (*74*). Data S6 shows results with TM-scores not less than 0.5.

## Acknowledgments

We thank members of the BLAST/STIM community for many helpful discussions on PAS domains in various model organisms.

## Author contributions

Conceptualization: JX, IBZ

Methodology: JX, VMG

Investigation: JX, VMG, IBZ

Supervision: IBZ

Writing—original draft: JX

Writing—review & editing: VMG, IBZ

## Data and materials availability

All data are available in the main text or the supplementary materials. Sequences, structure models, and sequence logos for clusters, and Python scripts used for analyses are available in GitHub https://github.com/bioliners/PAS.

