## Supplementary figures and tables for "Origin and functional diversification of PAS domain, a ubiquitous intracellular sensor"

**This PDF file includes:**

Figs. S1 to S8

Tables S1 to S5

References (75 to 156)

**Other Supplementary Materials for this manuscript include the following:**

Datasets S1 to S9

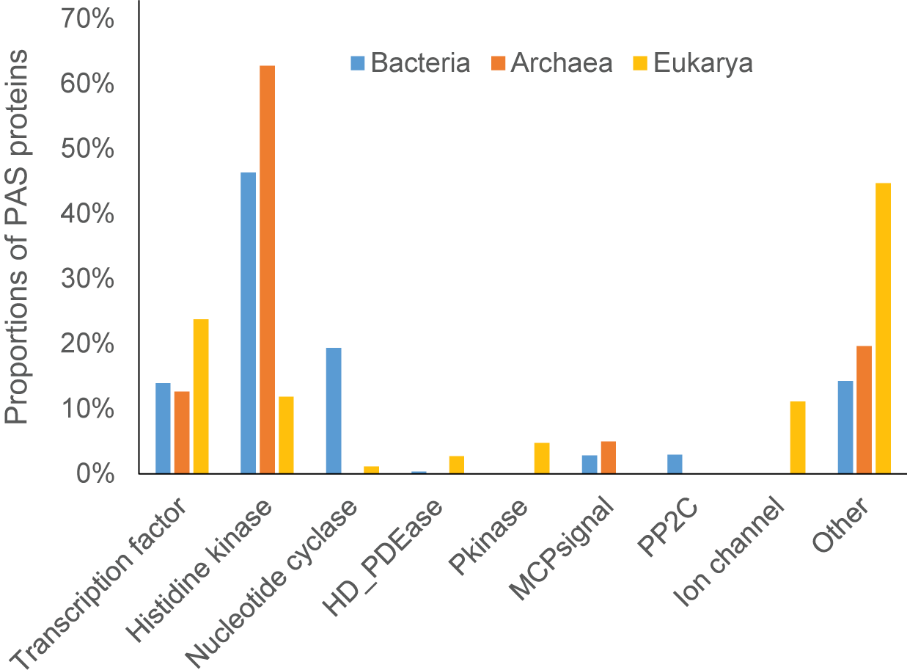

**Fig. S1. Different PAS containing proteins across bacteria, archaea, and eukaryotes.** Categories are defined by Pfam families. Transcription factor: HTH, HLH, GerE, GATA, Zn_clus, and Trans_reg_C; Histidine kinase: HisKA, HATPase_c, HWE, and His_kinase; Nuleotide cyclase: GGDEF, EAL, and Guanylate_cyc; HD_PDEase: HD and PDEase; Pkinase: Pkinase and PK_Tyr_Ser-Thr; MCPsignal: MCPsignal; PP2C: SpoIIE and PP2C; Ion channel: ion_trans.

**A**

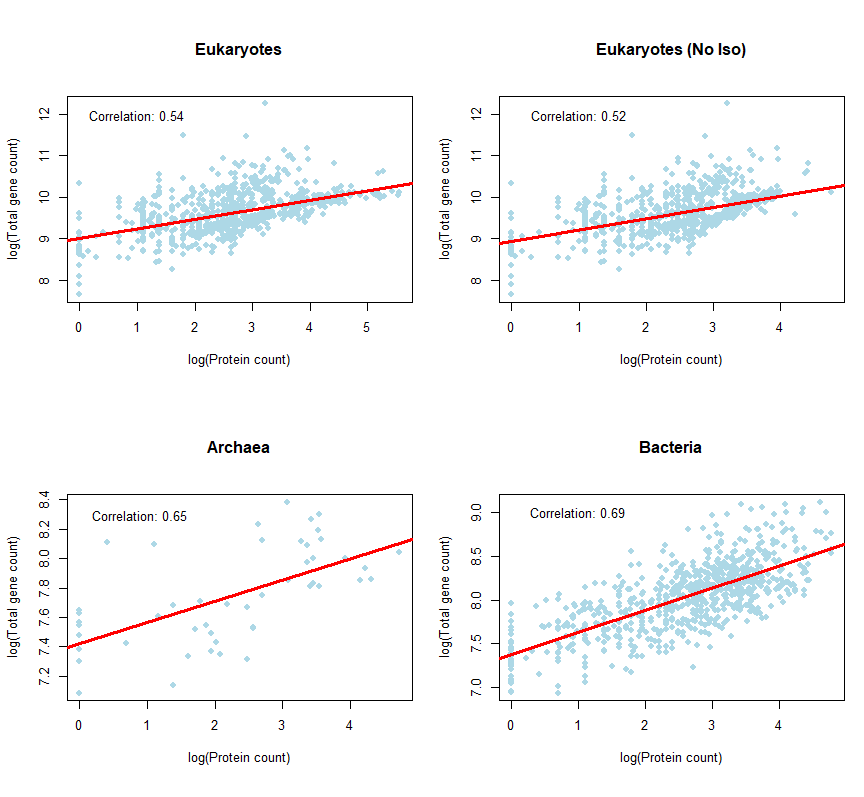

**B**

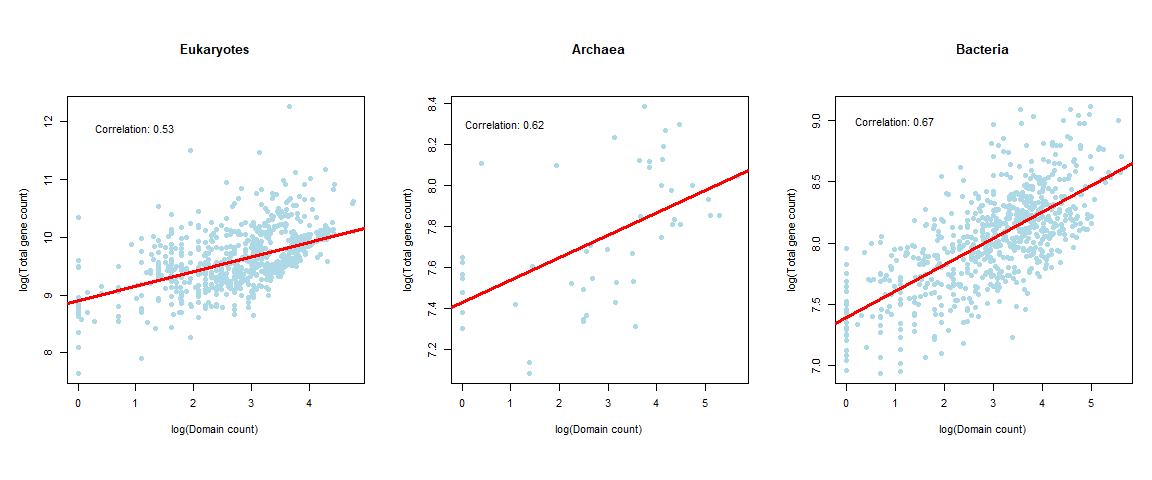

**Fig. S2. Spearman correlation of PAS proteins (A) and PAS domains (B) with the total number of genes.** Counts were normalized by total gene numbers at the family level (see Materials and Methods). Double-logarithmic scale is used. “No iso” indicates protein counts after removing isoforms. **A**. Eukaryotes: rho = 0.54, p-value < 2.2e-16, Eukaryotes (no iso): rho = 0.52, p-value < 2.2e-16, Archaea: rho = 0.65, p-value = 7.333e-07, Bacteria: rho = 0.69, p-value < 2.2e-16. **B**. Eukaryotes: rho = 0.53, p-value < 2.2e-16, Archaea: rho = 0.62, p-value = 2.871e-06, Bacteria: rho = 0.67, p-value < 2.2e-16. In the case of eukaryotes, the longest isoforms were used to illustrate the correlation of PAS domains with the total number of genes.

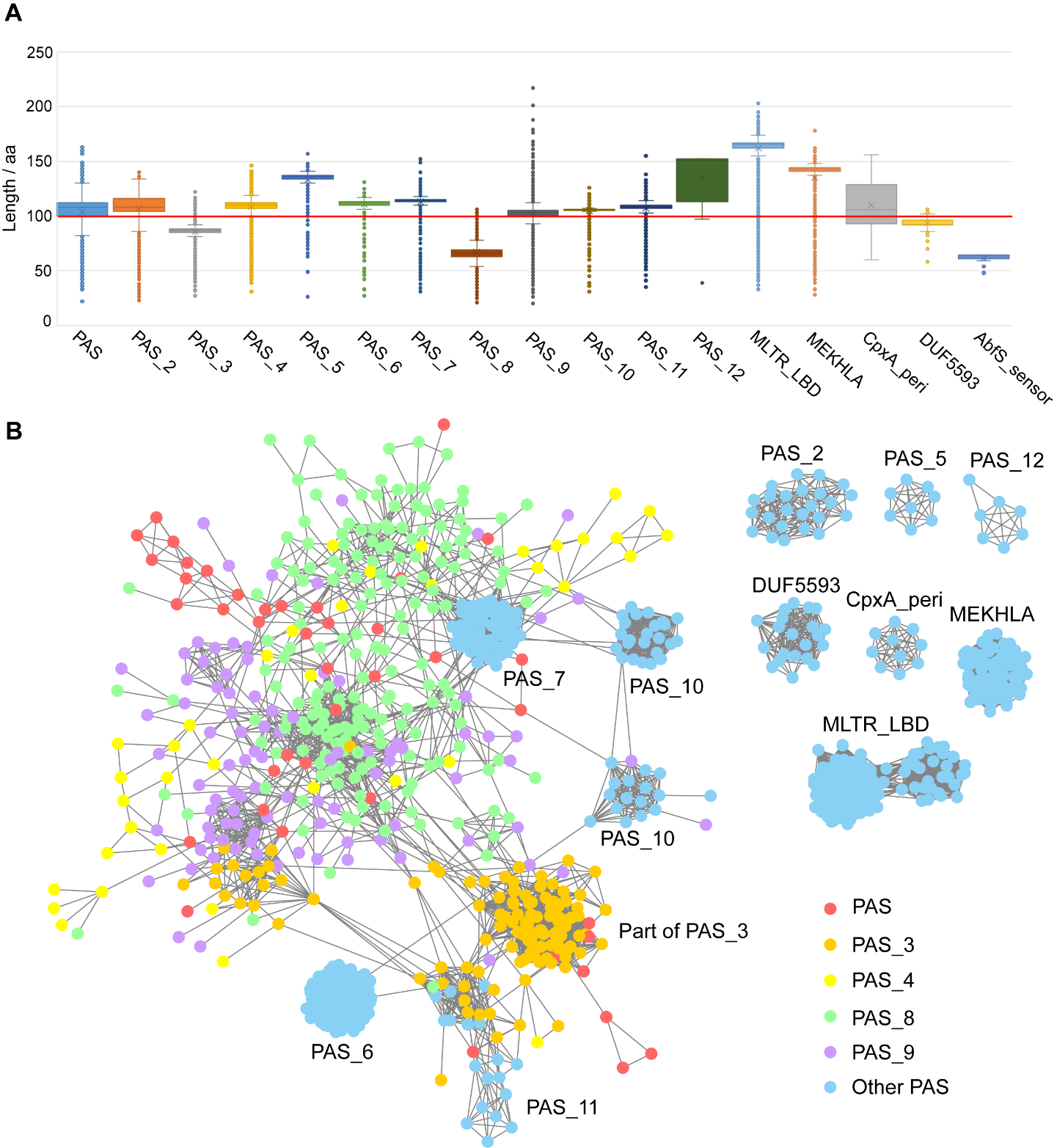

Fig. S3. PAS domain families in the Pfam database. (A) Sequence lengths of PAS families. The length of 100 amino acids is labeled by a red line. Sequences from PAS_3, PAS_8, and AbfS_sensor are substantially shorter than normal PAS domains. (B) Sequence similarity network of seed sequences. Mutual BLAST hits (E-value < 0.05, query coverage > 80%) are shown in Cytoscape. Blue nodes show separated families; other colors show overlapped families. Outlier sequences are not shown for simplicity.

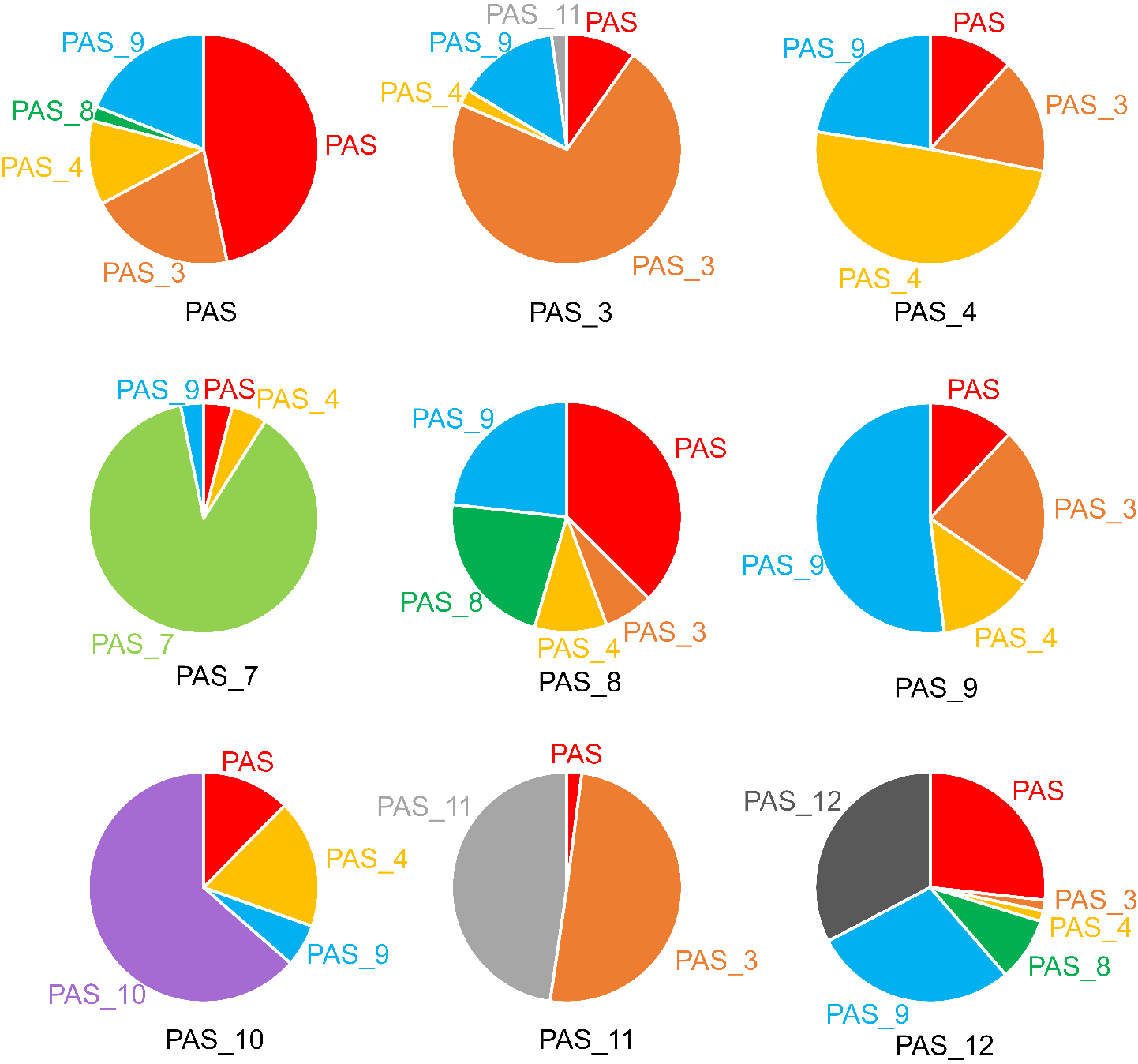

Fig. S4. Overlaps among Pfam PAS domain families. PAS domain sequences collected from RefSeq were searched against all Pfam families using *hmmscan*. Charts show the best matched HMM for sequences from each PAS family. The query HMM used for sequence collection are labeled below each chart. The rest of PAS families have few overlaps and are therefore not shown.

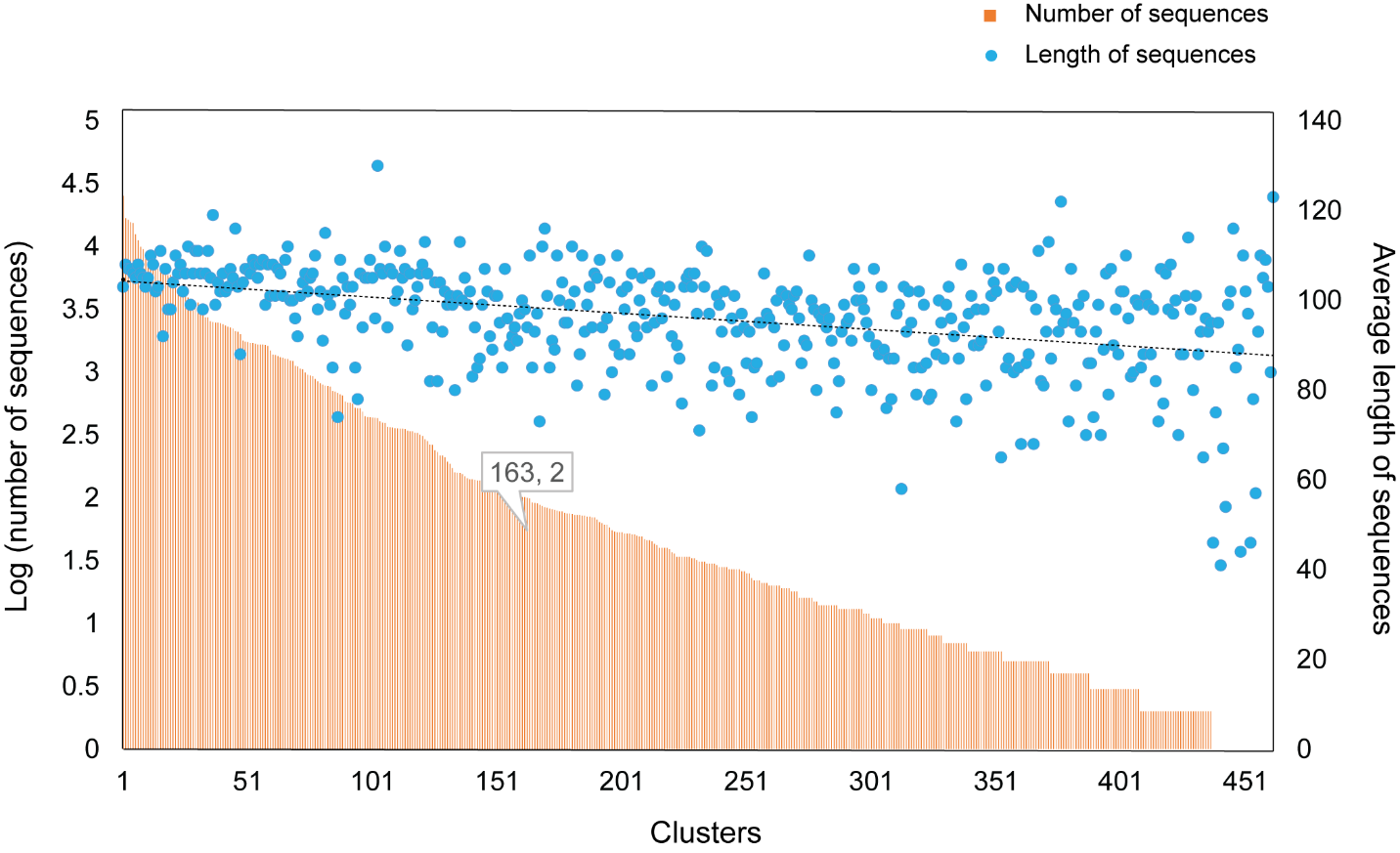

**Fig. S5. Size and sequence length of PAS domain clusters.** Clusters 1-462 were ordered by their numbers of PAS domain sequences from large to small. Yellow bars show the number of sequences in each cluster in the 10 based logarithm. Blue dots show the average length of sequences in each cluster. Clusters 1-163 contain at least 100 sequences, which is labeled in the plot.

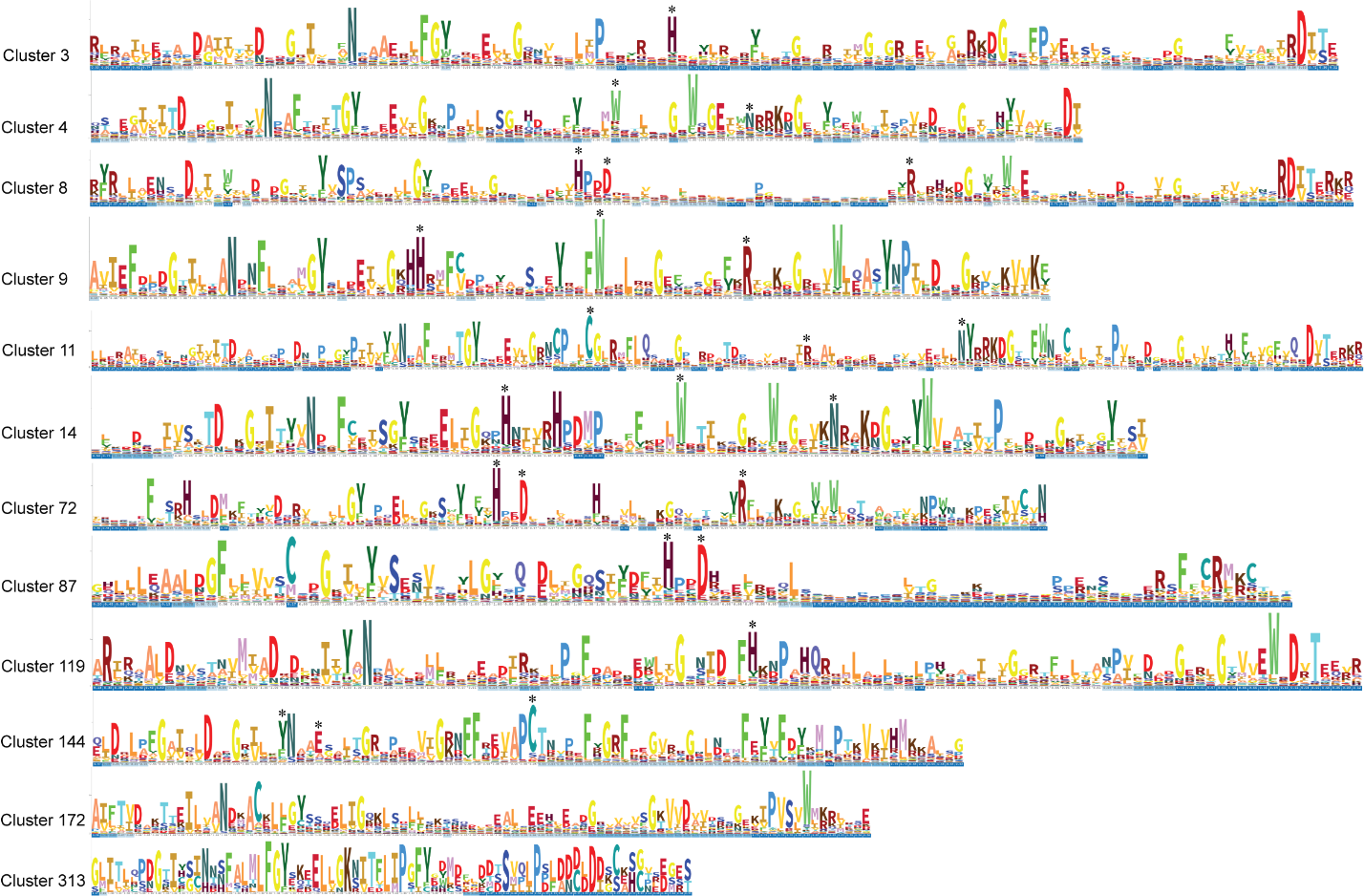

Fig. S6. Sequence logos of selected clusters. Gaps were trimmed by trimAl. Logos were built by Skylign. Conserved key residues are labeled with asterisks.

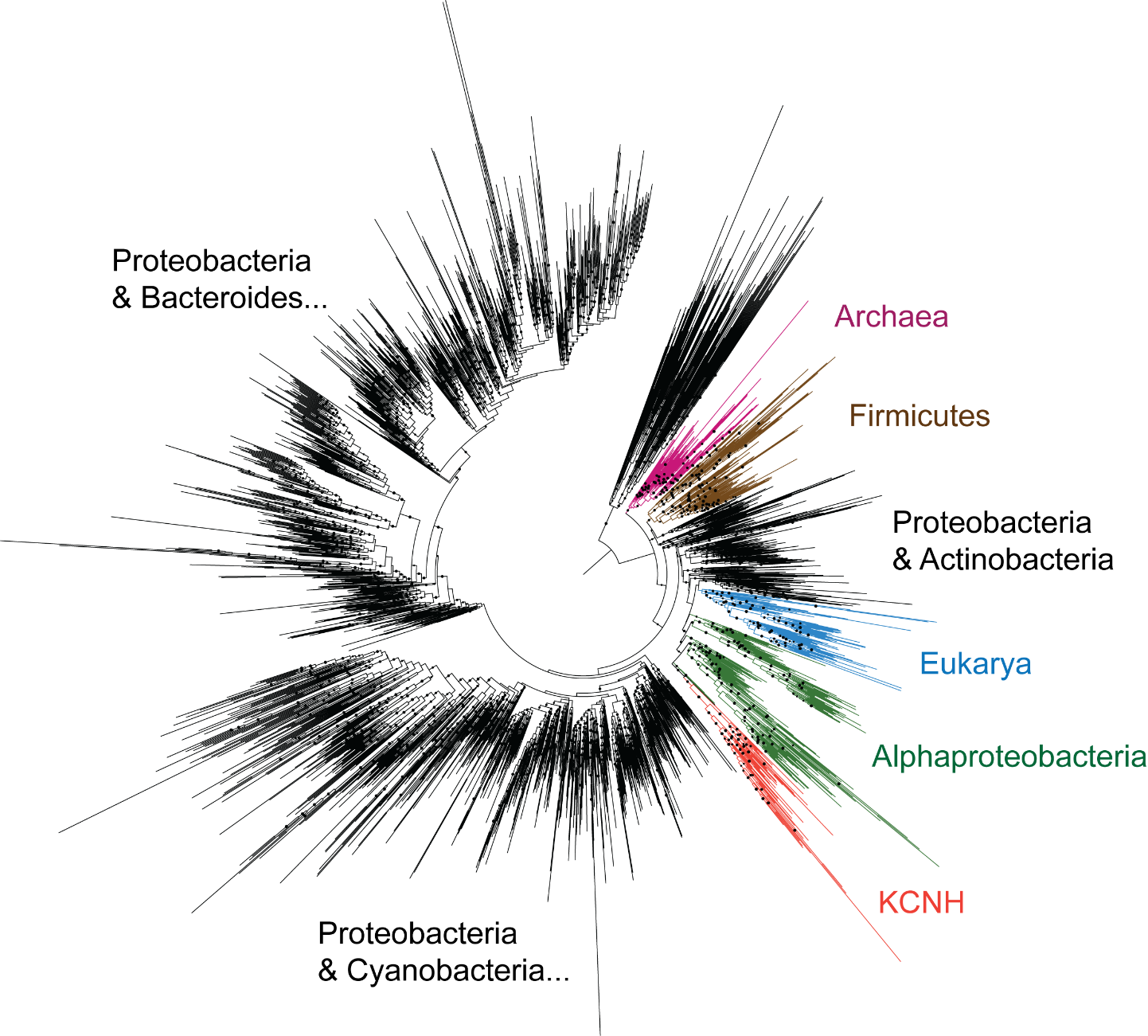

Fig. S7. Phylogenetic tree of cluster 11 (FMN). Sequences from cluster 11 were reduced at 60% redundancy. The maximum likelihood tree was built using LG+G4 model. Dots show bootstrap values > 70. Taxonomy was highlighted and labeled beside the tree. KCNH-PAS is close to PAS from Alphaproteobacteria.

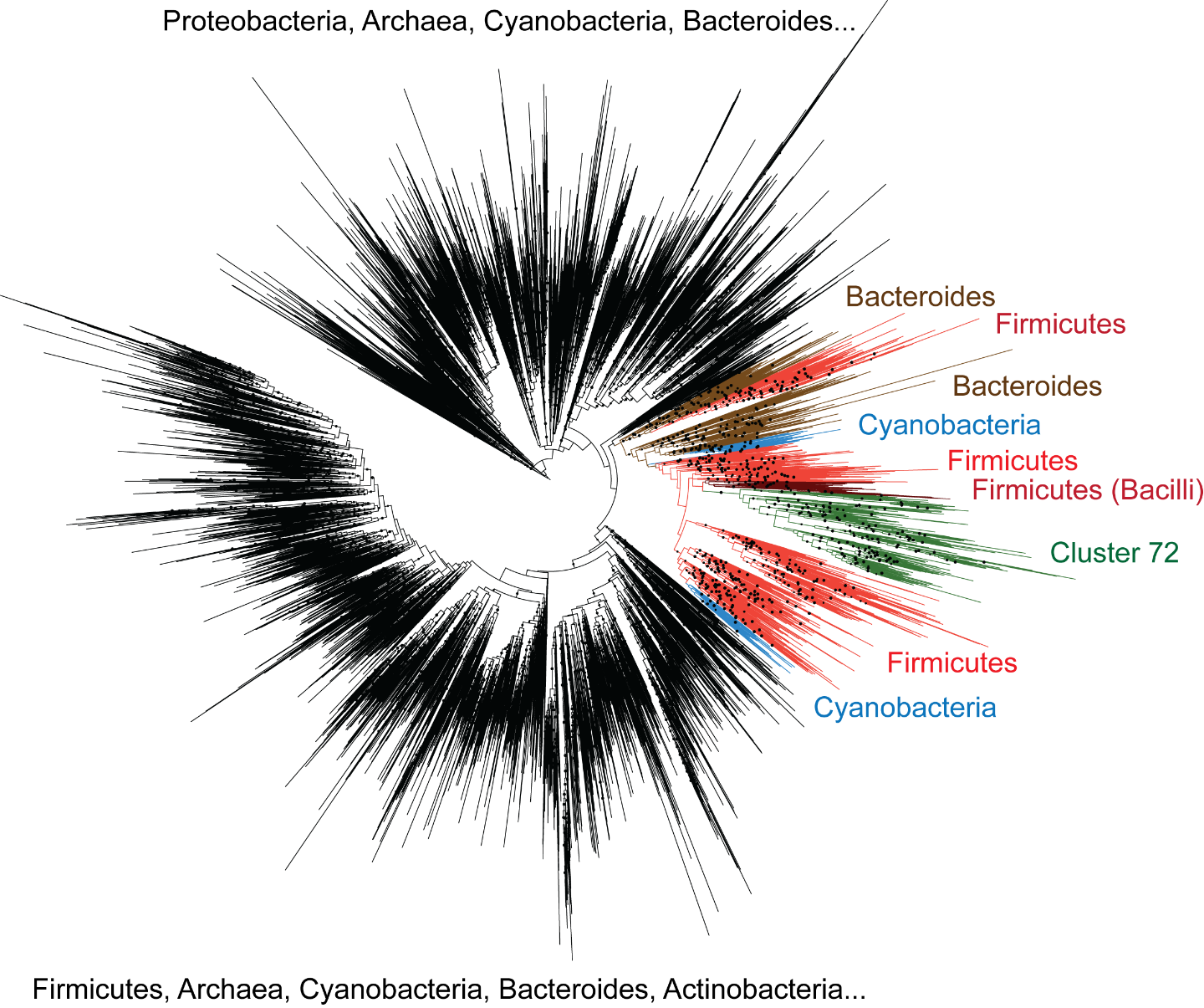

Fig. S8. Phylogenetic tree of clusters 8 and 72 (bHLH-PAS B). Sequences from cluster 8 and 72 were reduced at 60% redundancy. The maximum likelihood tree was built using LG+G4 model. Dots show bootstrap values > 70. Taxonomy was highlighted and labeled beside the tree. Cluster 72 is close to PAS from Firmicutes (Bacilli).

**Table S1 Cofactor-binding PAS domains**

| **Cofactors** | **Proteins** | **Example organisms** | **Pfam** | **PDB** | **References** |
| --- | --- | --- | --- | --- | --- |
| Heme | FixL | *Bradyrhizobium japonicum* | PAS | 1DRM | (*24*) |
|  | DosP | *Escherichia coli* | PAS_9 | 1V9Y | (*25*) |
|  | Aer2 | *Pseudomonas aeruginosa* | PAS_8 | 4HI4 | (*26, 75*) |
|  | CLOCK | *Homo sapiens* | PAS | - | (*39*) |
|  | NPAS2 | *Homo sapiens* | PAS | - | (*40*) |
|  | ERG3 | *Homo sapiens* | PAS_9 | - | (*38*) |
|  | RcoM | *Burkholderia xenovorans* | PAS_4? | - | (*76*) |
|  | PDEA1 | *Acetobacter xylinum* | PAS_9 | - | (*77*) |
|  | YybT | *Bacillus subtilis* | - | - | (*78*) |
|  | NtrY | *Brucella abortus* | PAS | - | (*79*) |
|  | Hpk2 | *Treponema denticola* | PAS_4 | - | (*80*) |
|  | SenX3 | *Mycobacterium tuberculosis* | PAS_4? | - | (*81*) |
|  | PpsR | *Rhodobacter sphaeroides* | PAS_9 | - | (*82*) |
|  | BdlA | *Pseudomonas aeruginosa* | PAS_9 | - | (*83*) |
|  | IcpB | *Azorhizobium caulinodans* | PAS_9 | - | (*84*) |
|  | Per2 | *Mus musculus* | PAS, PAS_3 | - | (*85-87*) |
| FAD | Aer | *Escherichia coli* | PAS_3 | 8DIK | (*29, 88*) |
|  | CetB | *Campylobacter jejuni* | PAS_3 | - | (*89*) |
|  | MmoS | *Methylococcus capsulatus* | PAS_9 | 3EWK | (*28*) |
|  | NifL | *Azotobacter vinelandii* | PAS | 2GJ3 | (*27*) |
|  | Vivid | *Neurospora crassa* | PAS_9 | 2PDR | (*90*) |
|  | WC-1 | *Neurospora crassa* | PAS_9 | - | (*7, 91*) |
|  | AerC | *Azospirillum brasilense* | PAS_9, PAS_3 | - | (*30*) |
| FMN | YtvA | *Bacillus subtilis* | PAS_9 | 2PR6 | (*92*) |
|  | phototropins | *Arabidopsis thaliana* | PAS_9 | 2Z6C | (*93*) |
|  | phy3 | *Adiantum capillus-veneris* | PAS_9 | 1G28 | (*94*) |
|  | LOV | *Rhodobacter sphaeroides* | PAS_9 | 4HIA | (*95*) |
|  | PpSB-LOV | *Pseudomonas putida* | PAS_9 | 5J4E | (*96*) |
|  | DsLOV | *Dinoroseobacter shibae* | PAS_9 | 4KUO | (*97*) |
|  | LovK | *Caulobacter cresentus* | PAS_9 | - | (*98*) |
|  | Lovhk | *Brucella abortus* | PAS_9 | 5EPV | (*99*) |
|  | EL222 | *Erythrobacter litoralis* | PAS_9 | 3P7N | (*100*) |
|  | Cagg_3753 | *Chloroflexus aggregans* | PAS_3 | 6RHG | (*101*) |
|  | aureochrome | *Phaeodactylum tricornutum* | PAS_9 | 5DKK | (*102*) |
|  | PAL | *Nakamurella multipartita* | PAS_9 | 6HMJ | (*103*) |
|  | ENV1 | *Trichoderma reesei* | PAS_9 | 4WUJ | (*104*) |
| *p*-coumaric acid | PYP | *Halorhodospira halophila* | PAS | 2PHY | (*34*) |
|  | Ppr | *Rhodospirillum centenum* | PAS? | 1MZU | (*105, 106*) |
|  | Ppd | *Thermochromatium tepidum* | PAS | - | (*107*) |
| Riboflavin | EL346 | *Erythrobacter litoralis* | PAS_9 | 4R38 | (*33*) |
| Fe-S cluster | NreB | *Staphylococcus carnosus* | - | - | (*47*) |

Table S2. Distribution of Pfam PAS families in Bacteria, Archaea, and Eukaryotes

| **PAS family** | **Bacteria** | **Archaea** | **Eukaryotes** |
| --- | --- | --- | --- |
| PAS | Yes | Yes | Yes |
| PAS_2 | Yes | No | Yes |
| PAS_3 | Yes | Yes | Yes |
| PAS_4 | Yes | Yes | Yes |
| PAS_5 | Yes | No | Yes |
| PAS_6 | Yes | No | Yes |
| PAS_7 | Yes | Yes | No |
| PAS_8 | Yes | Yes | Yes |
| PAS_9 | Yes | Yes | Yes |
| PAS_10 | Yes | Yes | No |
| PAS_11 | Yes | No | Yes |
| PAS_12 | Yes | No | No |
| MLTR_LBD | Yes | No | No |
| MEKHLA | Yes | No | Yes |
| DUF5593 | Yes | No | No |
| CpxA_peri | Yes | No | No |
| AbfS_sensor | Yes | No | No |

Table S3. PAS secondary structure elements in Pfam PAS HMMs.

| **PAS family** | **Secondary structures** | | | | | | | | | | | | | | | |
| --- | --- | --- | --- | --- | --- | --- | --- | --- | --- | --- | --- | --- | --- | --- | --- | --- |
| PAS |  |  | A’α | Aß | Bß |  | Cα | Dα | Eα | Fα |  |  | Gß | Hß | Iß |  |
| PAS_2 |  |  | A’α | Aß | Bß |  | Cα | Dα | Eα | Fα |  |  | Gß | Hß | Iß |  |
| PAS_3 |  |  |  | * | Bß |  | Cα | Dα | Eα | Fα |  |  | Gß | Hß | Iß |  |
| PAS_4 |  |  |  | Aß | Bß |  | Cα | Dα | Eα | Fα |  |  | Gß | Hß | Iß | Jα |
| PAS_5 | A’’’α | A’’α | A’α | Aß | Bß |  | Cα |  | Eα | Fα |  |  | Gß | Hß | Iß |  |
| PAS_6 |  |  | A’α | Aß | Bß |  |  |  |  | Fα |  |  | Gß | Hß | Iß | Jα |
| PAS_7 |  |  |  | Aß | Bß |  | Cα | Dα | Eα | Fα |  |  | Gß | Hß | Iß | Jα |
| PAS_8 |  |  | A’α | Aß | Bß |  | Cα | Dα | Eα | Fα |  |  | * | * | * |  |
| PAS_9 |  |  |  | Aß | Bß |  | Cα | Dα | Eα | Fα |  |  | Gß | Hß | Iß |  |
| PAS_10 |  |  | A’α | Aß | Bß |  | Cα | Dα | Eα | Fα |  |  | Gß | Hß | Iß |  |
| PAS_11 |  |  |  | Aß | Bß |  | Cα | Dα | Eα | Fα |  |  | Gß | Hß | Iß | Jα |
| PAS_12 | A’’’α | A’’α | A’α | Aß | Bß |  | Cα | Dα | Eα | Fα |  |  | Gß | Hß | Iß |  |
| MLTR_LBD |  |  | A’α | Aß | Bß |  | Cα | Dα | Eα | Fα | F’α | F’’α | Gß | Hß | Iß | Jα |
| MEKHLA |  | A’’α | A’α | Aß |  |  | Cα | Dα | Eα | Fα |  |  | Gß | Hß | Iß |  |
| DUF5593 |  |  |  | Aß | Bß | Cß |  |  | Eα | Fα |  |  | Gß | Hß | Iß |  |
| CpxA_peri** |  | A’’α | A’α | Aß | Bß |  |  |  |  | Fα |  |  | Gß | Hß | Iß | Jα |
| AbfS_sensor*** |  | A’’α | A’α | Aß |  |  | Cα | Dα | Eα |  |  |  | * | * | * |  |

PAS domains have a common structural fold: Aß-Bß-Cα-Dα-Eα-Fα-Gß-Hß-Iß. The representative sequence of each Pfam PAS family was identified using *hmmsearch* against seed sequences. Secondary structures for representative sequences were identified using Quick2D (https://toolkit.tuebingen.mpg.de/tools/quick2d).

*Essential regions in PAS_3, PAS_8, and AbfS_sensor were not covered by Pfam HMMs.

**Two extracellular Cache domains were mistakenly included in the PAS superfamily.

Table S4. Overlaps among Pfma PAS domain families

| **PAS family** | **RefSeq sequences** | **Correct sequences** | **Percentage of correctness** |
| --- | --- | --- | --- |
| PAS* | 648,279 | 300,617 | 46% |
| PAS_2 | 11,734 | 11,734 | 100% |
| PAS_3* | 481,799 | 344,493 | 72% |
| PAS_4* | 641,254 | 313,073 | 49% |
| PAS_5 | 4,049 | 4,049 | 100% |
| PAS_6 | 11,407 | 11,402 | 100% |
| PAS_7* | 64,296 | 55,951 | 87% |
| PAS_8* | 241,387 | 52,895 | 22% |
| PAS_9* | 577,171 | 299,371 | 52% |
| PAS_10* | 23,131 | 14,674 | 63% |
| PAS_11* | 35,694 | 16,956 | 48% |
| PAS_12* | 269 | 88 | 33% |
| MLTR_LBD | 58,261 | 58,200 | 100% |
| MEKHLA | 3,572 | 3,262 | 91% |
| DUF5593 | 1, 524 | 1,524 | 100% |
| CpxA_peri | 2,575 | 2,575 | 100% |
| AbfS_sensor | 93 | 93 | 100% |

Sequences for each PAS family were collected from RefSeq by *hmmsearch* using Pfam HMMs as queries (E-value < 0.01). For some PAS families, query HMMs also matched sequences from other families. Correct sequences are defined as sequences with the best match to the query HMM using *hmmscan* (Fig. S4).

*Nine PAS families have overlapped sequences.

**Table S5 PAS proteins in the human proteome.**

| **Class** | **Protein** | **Function** | **Associated Disease** | **Ref** |
| --- | --- | --- | --- | --- |
| KCNH | ERG1 (KCNH2) | Heart rhythm | Long/short QT syndrome | (*108, 109*) |
|  | ERG2 (KCNH6) | Expressed in CNS |  | (*110*) |
|  | ERG3 (KCNH7) |  |  | (*110*) |
|  | EAG1 (KCNH1) |  | Cancer, Temple-Barrister syndrome, Zimmerman-Laband syndrome | (*111-113*) |
|  | EAG2 (KCNH5) |  | Cancer | (*111, 114*) |
|  | ELK1 (KCNH4) |  |  | (*115*) |
|  | ELK2 (KCNH3) |  |  | (*115*) |
|  | ELK3 (KCNH8) |  |  | (*116*) |
| PDE8 | PDE8A | cAMP hydrolyzation | Depression | (*53, 117*) |
|  | PDE8B |  | Adrenal hyperplasia | (*118*) |
| bHLH-PAS | AHR | Ligand sensing | Cancer, immune diseases | (*11, 119, 120*) |
|  | AHRR | AHR repressor | Cancer | (*121*) |
|  | HIF1α | Hypoxia responses | Cancer, cardiovascular diseases | (*8, 10, 122*) |
|  | HIF2α (EPAS1) |  | Cancer, erythrocytosis | (*12, 123, 124*) |
|  | HIF3α (IPAS) | HIF repressor | Cancer, Parkinson’s disease | (*125-127*) |
|  | SIM1 | Neural development | Obesity | (*128*) |
|  | SIM2 |  | Cancer, Down syndrome | (*129-131*) |
|  | NPAS1 |  | Psychiatric diseases | (*132, 133*) |
|  | NPAS3 |  | Schizophrenia | (*134, 135*) |
|  | NPAS4 | Neural development, DNA repair | Schizophrenia, diabetes, aging | (*136-138*) |
|  | CLOCK | Circadian rhythm | Circadian disorders | (*139*) |
|  | NPAS2 |  | Circadian disorders | (*140*) |
|  | PASD1 | CLOCK repressor, cancer testis antigen | Circadian disorders, multiple myeloma, lymphoma | (*141-143*) |
|  | ARNT | bHLH dimerization | Cancer | (*144*) |
|  | ARNT2 |  | Cancer | (*144*) |
|  | BMAL1 (ARNTL) |  | Circadian disorders | (*145*) |
|  | BMAL2 (ARNTL2) |  | Circadian disorders | (*146*) |
|  | NCOA1 (SRC1) | Transcriptional coactivator | Cancer | (*147, 148*) |
|  | NCOA2 (SRC2) |  | Metabolic diseases | (*149*) |
|  | NCOA3 (SRC3) |  | Cancer, hearing loss | (*150, 151*) |
|  | PER1* | Circadian rhythm | Circadian disorders, long-term memory loss, hypertension | (*152, 153*) |
|  | PER2* |  | Circadian disorders, cardiovascular diseases | (*154*) |
|  | PER3* |  | Circadian disorders, mood disorders | (*155*) |
| PASK | PASK | Signal transduction | Metabolic diseases | (*156*) |

* PER1-3 do not have bHLH domains but function together with bHLH-PAS proteins.

Dataset S1. Presence of PAS domains in UniProt reference proteomes.

Data contains 22,925 reference proteomes from UniProt. PAS domain presence in each proteome is shown.

Dataset S2. PAS domain containing proteins identified in InterPro.

Data contains all PAS-containing proteins from InterPro defined by the Pfam PAS fold.

Dataset S3. PAS domains identified in 14 representative eukaryotic genomes.

Data contains a summary sheet and 14 sheets with PAS-containing proteins in each genome.

Clusters to which PAS domains belong are shown in the file. PAS domains that belong to well-defined clusters and have conserved key residues are highlighted (blue, heme-binding; red, flavin-binding). Isoforms are highlighted in red.

Dataset S4. PAS domains in 43 representative eukaryotic genomes.

Data contains a summary sheet and 43 sheets with PAS-containing proteins in each genome. Isoforms and false positive hits are highlighted in red.

Dataset S5. Sequences of PAS domains identified in the RefSeq database.

Data contains all PAS domain sequences used for clustering in FASTA format. Missed regions from PAS_3 and PAS_8 sequences were added (20 amino acids at N-terminal PAS_3 and 50 amino acids at C-terminal PAS_8). Sequences for PAS, PAS_3, PAS_4, PAS_7, PAS_8, PAS_9, PAS_10, PAS_11, and PAS_12 were combined and reduced 80% sequence redundancy using CD-HIT.

Dataset S6. RefSeq accession numbers for proteins with PAS domains assigned to MCL clusters.

Each column contains PAS domains of a cluster.

Dataset S7. Phyletic distribution and domain architecture of PAS domain containing proteins in the RefSeq database.

Each sheet contains PAS-containing proteins from a well-defined cluster with the conserved key residues for cofactor-binding.

Dataset S8. PAS domains in the human genome identified by the structure search.

Data contains human proteins matching to 2KDK (TMscore > 0.5).

Dataset S9. Results of BLAST searches with representative eukaryotic PAS domains.

Each sheet contains the top 100 BLAST hits using a eukaryotic PAS domain as query (results are ordered by sequence identities). The top BLAST hits from bacterial proteins are highlighted in yellow.
